# Pro-inflammatory cytokines drive deregulation of potassium channel expression in primary synovial fibroblasts

**DOI:** 10.1101/2020.02.06.937078

**Authors:** Omar Haidar, Nathanael O’Neill, Caroline A. Staunton, Selvan Bavan, Fiona O’Brien, Sarah Zouggari, Umar Sharif, Ali Mobasheri, Kosuke Kumagai, Richard Barrett-Jolley

## Abstract

The synovium secretes synovial fluid, but is also richly innervated with nociceptors and acts as a gateway between avascular joint tissues and the circulatory system. Resident fibroblast-like synoviocytes’ (FLS) calcium-activated potassium channels (K_Ca_) change in activity in arthritis models and this correlates with FLS *activation*.

**Objective:** To investigate this activation in an *in vitro* model of inflammatory arthritis; 72hr treatment with cytokines TNFα and IL1β.

**Methods:** FLS cells were isolated from rat synovial membranes. We analysed global changes in FLS mRNA by RNA-sequencing and then focused on FLS ion channels genes and corresponding FLS electrophysiological phenotype, finally modelling data with Ingenuity Pathway Analysis (IPA) and MATLAB.

**Results:** IPA showed significant activation of inflammatory, osteoarthritic and calcium signalling canonical pathways by cytokines, and we identified ~200 channel gene transcripts. The large K_Ca_ (BK) channel consists of the pore forming Kcnma1 together with β-subunits. Following cytokine treatment, a significant increase in Kcnma1 RNA abundance was detected by qPCR and changes in several ion channels were detected by RNA-sequencing, including a loss of BK channel β-subunit expression Kcnmb1/2 and increase in Kcnmb3. In electrophysiological experiments, there was a decrease in overall current density at 20mV without change in chord conductance at this potential.

**Conclusion:** TNFα and IL1β treatment of FLS in vitro recapitulated several common features of inflammatory arthritis at the transcriptomic level, including increase in Kcnma1 and Kcnmb3 gene expression.

## Introduction

Rheumatoid arthritis (RA) and osteoarthritis (OA) are degenerative diseases that target articular joint structures resulting in pain, loss of function and frequent disability. Whilst RA is an established inflammatory condition, the contribution of inflammatory processes to OA was less well known until recently. Mediators of inflammation (i.e. including pro-inflammatory cytokines) contribute to the development of synovitis, which is known to drive disease progression in RA. In contrast, it is thought that in OA, synovitis can be caused by the release of cartilage fragments and meniscal damage that in turn, activate synovial lining cells (Fernandez-Madrid *et al.* 1995; Roemer *et al.* 2013; Mathiessen and Conaghan 2017). How inflammation drives joint destruction is not fully known. One feature of synovitis, however, is the presence of major pro-inflammatory cytokines, such as Interleukin-1β (IL-1β), tumour necrosis factor alpha (TNFα) and interleukin 6 (IL-6), resulting in suppression of collagen and proteoglycan synthesis, increased inflammatory signalling, and protease expression and activation (Martel-Pelletier *et al.* 1999).

Management of arthritis has significantly improved in recent years’, however, remission is rarely achieved, and many patients remain unresponsive to conventional and/or biologic treatments. In addition, current therapies and treatments are associated with notable side effects that can pose great challenges for long-term treatment, especially in patients with cardiovascular co-morbidities. For example, some drugs can increase the risk of cardiovascular disease or significantly impair immune responses, rendering patients more susceptible to infections and cancer (Kahlenberg and Fox 2011). Therefore, new therapeutic options and novel targets are needed that lead to pronounced improvement without inducing unwanted side effects and thus avoiding the need for joint replacement.

The synovium is the major barrier between the joint and the systemic circulation and plays a role in maintaining the health of articular cartilage (Sutton *et al.* 2009; Berenbaum 2013). The synovium lubricates the articular surfaces and provides nutrients for chondrocytes within the avascular cartilage; it has been suggested that catabolic enzymes such as matrix metalloproteinase (MMPs) are produced by synovial cells and diffuse into the cartilage. The intimal lining layer of the synovium produces lubricious synovial fluid and is composed of two cell types in relatively equal proportions: Type A or macrophage-like synovial cells and Type B or fibroblasts like synoviocytes (FLS). FLS cells contribute to the structural integrity of the joints by controlling the composition of the synovial fluid and extracellular matrix (ECM) of the joint. The synovial environment changes physically, chemically and physiologically with injury or the onset of disease and is thought to be a mediator in arthritis pain (Grubb 2004; Sellam and Berenbaum 2010; Kumahashi *et al.* 2011). FLS cells have been implicated in arthritis as they exhibit a transformed phenotype with increased invasiveness and production of various pro-inflammatory mediators that perpetrate inflammation and proteases that contribute to cartilage destruction (Noss and Brenner 2008; Bartok and Firestein 2010). Understanding the biology and regulation of FLS cells provides insight into the pathogenesis of inflammatory arthritis. FLS cells could potentially be targeted pharmacologically to produce increased volumes of synovial fluid as an alternative to intra-articular hyaluronan or synthetic fluid injection therapies. They are also a plausible analgesic target because they may interact with sensory neurons and have been described as “amplifiers” of neuropeptide mediated inflammation and pain.

To deepen our understanding of the synovium in the context of synovial joint health and disease, the electrophysiological profile of FLS cells needs to be characterised, along with the ion channels that are present. Ion channels are an essential component of any cell membrane that controls ion movement in and out of the cell and play an important role in a multitude of cell regulating processes, typically by modulating the membrane potential. Electrophysiological techniques have been used to characterise the biophysical properties of a number of different FLS preparations, including mouse, rabbit, bovine and human (Large *et al.* 2010; Friebel *et al.* 2014; Clark *et al.* 2017). The best available whole-cell mathematical model of the FLS is heavily dominated by a Ca^2+^-activated potassium conductance, with small added components of inward rectifiers, background and leak. A recent study by Kondo et al. demonstrated that human FLS express high levels of Ca^2+^-activated potassium channels and these ion channels were also identified in both rheumatoid arthritis-derived and rodent model FLS studies. Typically, Ca^2+^-activated channels couple with Ca^2+^ entry channels such as transient receptor potential (TRP) channels; they are both activated by the Ca^2+^ ions that enter and maintain the membrane potential hyperpolarised to “draw in” further Ca^2+^. In FLS, Ca^2+^-activated potassium channels appear to drive invasiveness of synoviocytes and progression of arthritis in both human and rodent RA models (Petho *et al.* 2016; Tanner *et al.* 2019), by increasing production of both inflammatory mediators and catabolic enzymes (Hu *et al.* 2012; Friebel *et al.* 2014; Tanner *et al.* 2015). This is a paradoxical effect for a potassium conductance, that would be predicted to hyperpolarise cells and reduce migration, proliferation and activity in general.

The synovium is an obvious target for the development of novel interventions in both RA and OA. The role of synovitis, the low-grade inflammation of the synovial lining of the joint, in OA progression is gradually emerging. Therefore, in this work we investigate the pathophysiology of cytokine induced synovitis in cultured synovial cells.

We investigate whether the TNFα and IL1β cytokine *in vitro* model of inflammation leads to a significant change of the BK ion channel and quantify the mechanism of this change. We use a combination of Next Generation RNA Sequencing (NGS), qPCR and patch-clamp electrophysiology to uncover several changes in potassium channel gene expression together with changes in cellular phenotype which involves a phenotypic switch in response to inflammation.

## Methods

### Further methodological details are included in the Supplementary Methods section

#### Animals

Synovial cells were prepared from tissue from rats euthanised by Home Office Approved methods for unassociated reasons in line with the ARRIVE Guidelines. All rats were untreated/wild-type male adult Wistar.

#### Preparation of FLS cells

Synovial fibroblasts were isolated from rat knee joints as described previously. Briefly, patella and menisci with attached synovial membrane were isolated and placed in 12-well plates in low glucose DMEM (Thermo Fischer - UK) with 20% foetal bovine serum (FBS), 100 U/ml penicillin, 100 µg/ml streptomycin and 2.5 µg/ml Amphotericin B (Thermo Fischer - UK) at 37 °C in a 5% CO2 incubator. For the first 7 days medium was replaced daily whilst out-growing FLS emerged from the tissues. At 7 days residual tissue was discarded and cells cultured as normal; when confluent, cells were detached from the flask surface by 1% Trypsin-EDTA solution. The suspension was centrifuged (340 x g, 5 min) and the resulting pellet was resuspended in culture medium (as above).

#### lmmunohistochemistry

In brief, cell suspension at a density of 2.5×104 cell/ml was transferred to multiwall plates and fixed with 2% paraformaldehyde in PBS at room temperature. CD248 (an FLS marker) immunohistochemistry was performed using rabbit anti-CD248 primary antibody (Ab, 1:100 dilution; AbCam, Cambridge) and FITC-conjugated donkey Anti-rabbit IgG secondary antibody (Ab, 1:500 dilution; Jackson ImmunoResearch Laboratories). Non-specific binding was blocked as previously described (Mobasheri *et al.* 2005). After 24 hr at 4 °C, cells were washed 3 times for 5 min with 0.05% SSC-20 and 0.005% Triton-X100. Slides were finally dipped into distilled water, air dried and mounted with mounting media (Vectashield with DAPI). Cells were visualised with confocal microscopy.

#### Real-Time PCR

RNA extraction was carried out using the RNeasy Plus Micro kit, together with gDNA eliminator and MinElute spin columns (Qiagen, UK). cDNA synthesis (mRNA) was performed using the RT2 First Strand kit (Qiagen, NL) according to the manufacturer’s protocol. qPCR analysis was performed using the Stratagene MX3000P RT-PCR System (Stratagene, La Jolla, CA) in a 25-µL reaction mixture. Expression relative to housekeeper Rplp1 was calculated as ΔCt.

#### RNA “Next Generation Sequencing” (NGS)

Figure 1 summarises our NGS workflow. In brief, cells were treated with 10 ng/ml of both TNF-α + IL1β (TNF-α from Thermo Fischer - UK and IL1β from R&D systems) for 72 hr. RNA extraction was carried out using the RNeasy Plus Micro kit (Qiagen, UK) according to the manufacturers protocol. Any RNA samples with concentrations of less than 5µg/ml and/or purity (260/280 and 260/230) less than 1.8 were excluded. RNA samples were sent to GATC-Biotech, Germany for sequencing, Samples were read as paired-end with a sequencing depth of 30M and a read length of 50bp. Further details are included in *Supplementary Methods.* Further bioinformatics were performed with R, for example the DPAC DA-PCA package or our local installation of the Galaxy server suite (Afgan *et al.* 2018). Specific packages are mentioned in the text, **Supplementary Methods** and Figure 1. **Full data are available on the EBI Array express database with Accession Number: E-MTAB-7798**.

**Figure 1.**
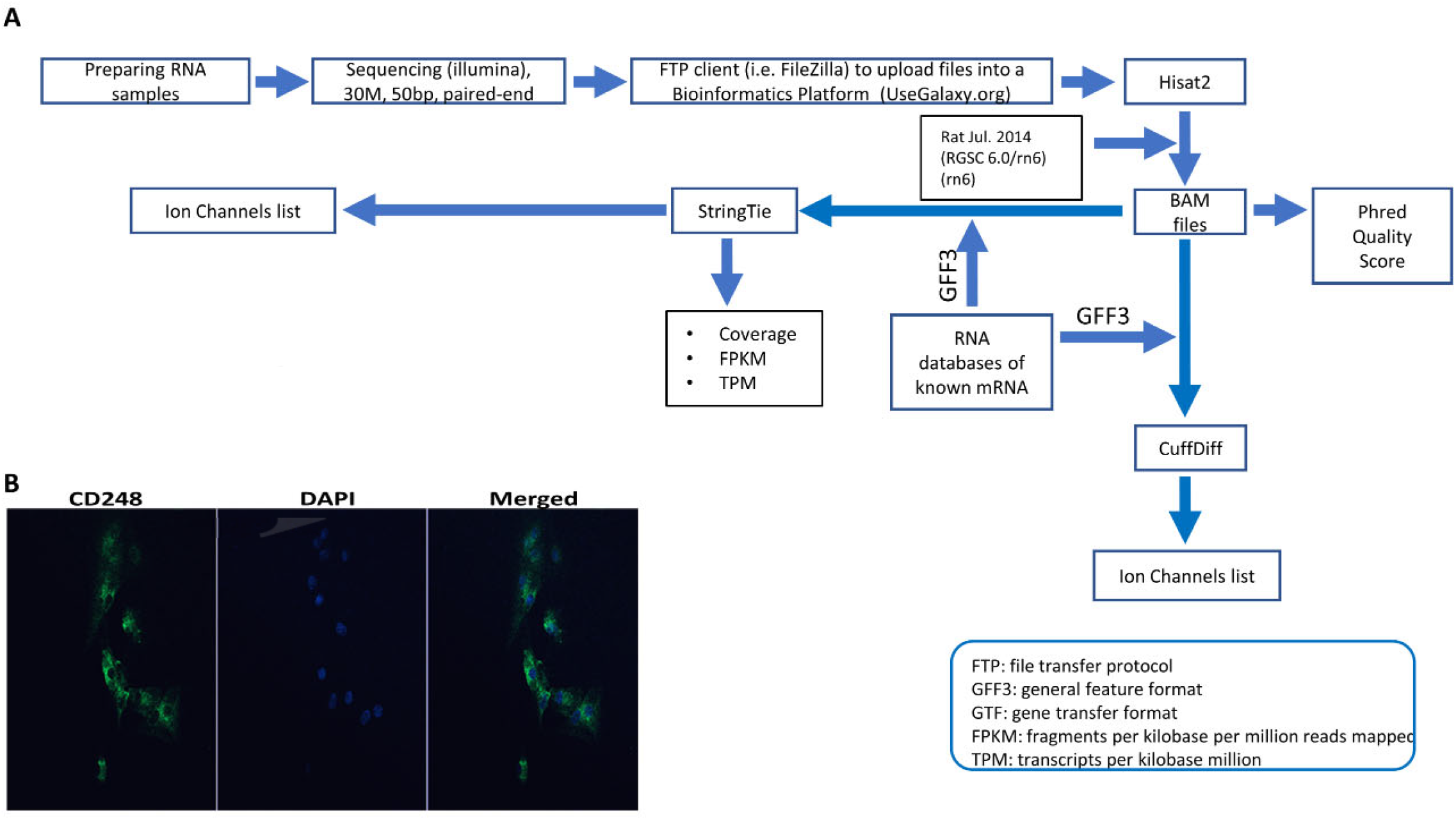
Experimental pipeline and verification. **(A)** Schematic of the Next Generation Sequencing (NGS) Pipeline. RNA was extracted from synovial tissue from 8 animals, were split into control and test groups and the test samples were treated with 10ng IL1β and TNFα as described in the *methods.* These 8 samples sequenced with Illumina and raw data files were uploaded using a FTP client to a bioinformatics platform for analysis. Within the bioinformatics platform, reads were aligned/mapped with Hisat2 program using the reference genome Rat Jul. 2014 (RGSC 6.0/rn6) (rn6). The resulting BAM files were used to measure phred quality score for all samples. Also, BAM files along with an annotation file (Rat GFF3) to assemble mapped reads and quantify gene expression. Such assembly and quantification took place with StringTie and CuffDiff package/tool that generated gene abundance estimates and differentially expressed genes, respectively. After that, using several servers and a local custom-script in MATLAB software allowed us to filter the transcriptome for all known ion channel genes for further analysis (for example Tables I and II). **(B)** Expression of CD248 in FLS cells. Immunofluorescence FLS cells showing the FLS marker CD248). DAPI was used for nuclear counterstaining (see *supplementary methods for details*).

#### Electrophysiology

Electrophysiology was performed as described previously (Lewis *et al.* 2013) but using isolated FLS. Intracellular solution was 115 mM Gluconic acid/Potassium salt, 26 mM KCl, 1 mM MgCl_2_ (BDH, VWR International Ltd), 5 mM Ethylene glycol tetraacetic acid (EGTA), 10 mM HEPES, pH 7.2. Extracellular (bath) solution was 140 mM NaCl, 5 mM KCl, 2 mM CaCl_2_ (Fluka Analytical cat#: 21114), 1 mM MgCl_2_, 10 mM HEPES, and 5 mM Glucose, osmolality approximately 300mOsm, pH 7.4. Junction potential - 14.4mV (Lewis *et al.* 2013). Thick-walled patch-pipettes were pulled from borosilicate glass capillary tubes (outer diameter 1.5 mm, inner diameter 0.86 mm; Intracel, UK) and gave a resistance of ~8 MΩ when filled. Whole cell patch-clamp electrophysiology was performed on FLS cells using a Cairn Optopatch amplifier (Cairn Research, UK). To compare voltage-gated currents, we performed whole-cell patch clamp experiments with voltage steps starting from a holding potential of −80mV for 2s (main text Figure 4). Recordings were filtered at 1kHz, digitized at 3kHz and recorded on a computer using WinWCP 5.3.4 software (John Dempster, Strathclyde University, UK). All experiments were performed at room temperature (18-22°C), and the results are expressed as the mean±SEM.

Analysis was performed using WinWCP 5.3.4 software (John Dempster, Strathclyde University, UK). Boltzmann curve fits were computed in MATLAB through nonlinear least squares optimization.

## Results

### Enriched pathways in FLS cells after cytokine treatment

RNA-seq detected 33251 transcripts and the **full data are available on the EBI Array express database with Accession Number: E-MTAB-7798.** The top (highest FPKM) expressed 10 genes were similar between control and cytokine treated cells (Table S1 and S2). Our first bioinformatic analysis tested the reproducibility of the 72hr, 10ng/ml TNFα + IL1β treatment. We used discriminant analyses of principle components with the DPAC package to show good separation of population (Figure 2), the genes primarily discriminating the treatment and control populations are largely those well established to be important for joint function, including several collagens and a Matrix metallopeptidase (MMP2). Ingenuity Pathway (IPA) Analysis (Qiagen, UK) was then used to identify the upstream regulators of the global differential expression pattern. This analysis predicted the top two regulators to be TNFα and IL1β (p-values 3e-17 and 6e-13 respectively), this is unsurprising since this was indeed the treatment regimen.

**Figure 2.**
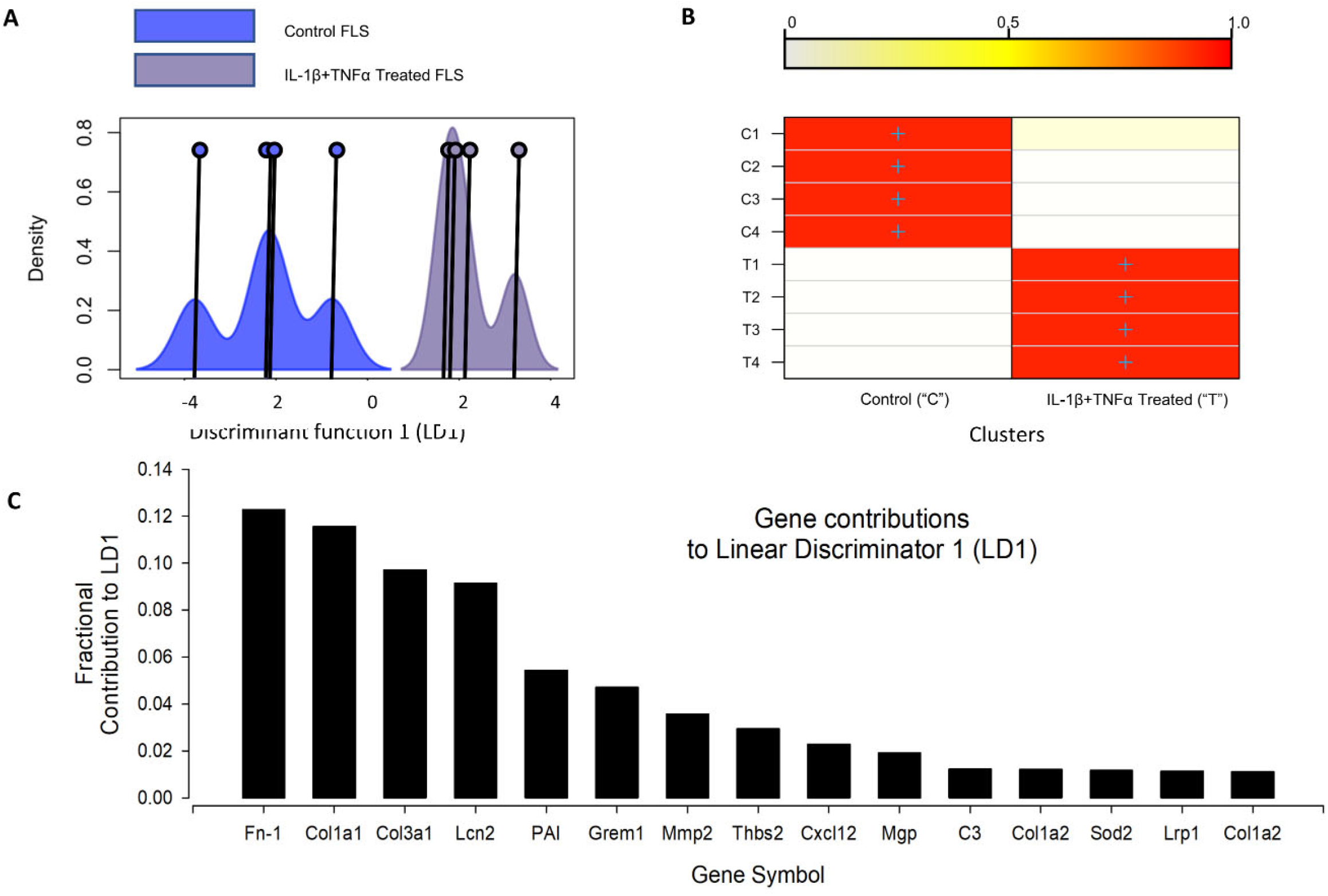
Discriminant Analyses of Global IL-1β TNFα Treatment Effects. **(A)** Shows the kernel density plots of the discriminant component co-ordinates for the control and IL-1β /TNFα treatment groups; co-ordinates on the x-axis and density on the y-axis; Individual co-ordinate centre points are illustrated by the vertical line and circle. There is clear separation between control and cytokine treated FLS samples. **(B)** A graphical confusion matrix showing the actual group membership (y-axis) and predicted cluster membership (on the y-axis). All groups are correctly clustered with greater than 0.9 probability. (C) The top 15 contributors to the linear discriminator function; Fn-1=fibronectin 1, Col1a1=collagen type I alpha 1 chain, Col3a1=collagen type III alpha 1 chain, Lcn2=lipocalin 2, PAI=Serpine1, Grem1=Gremlin 1, Mmp2=Matrix metallopeptidase 2, Thbs2=Thrombospondin 2, Cxcl12=C-X-C motif chemokine ligand 12, Mgp=Matrix Gla protein, C3=Complement C3, Col1a2=Collagen type I alpha 2 chain, Sod2=Superoxide dismutase 2, Lrp1=LDL receptor related protein 1, Col1a2=collagen, type I, alpha 2.

Figure 3 shows the canonical calcium signally pathway (p<0.5e-7), which was enriched following cytokine treatment. In addition, the rheumatoid (p<5e-13) and osteoarthritis (p<1e-9) pathways and cellular movement and proliferation canonical pathways (predicted *activation,* p<1e-13 in both cases) were also enriched following cytokine treatment (data not shown). In all cases, TNFα was determined to be the top *causal* agent, but four ion channels were also significant causal regulators (adjusted p<0.05) of the transcript-wide treatment changes; Clcn5, Trpv4, Trpv1, Kcnn4.

**Figure 3.**
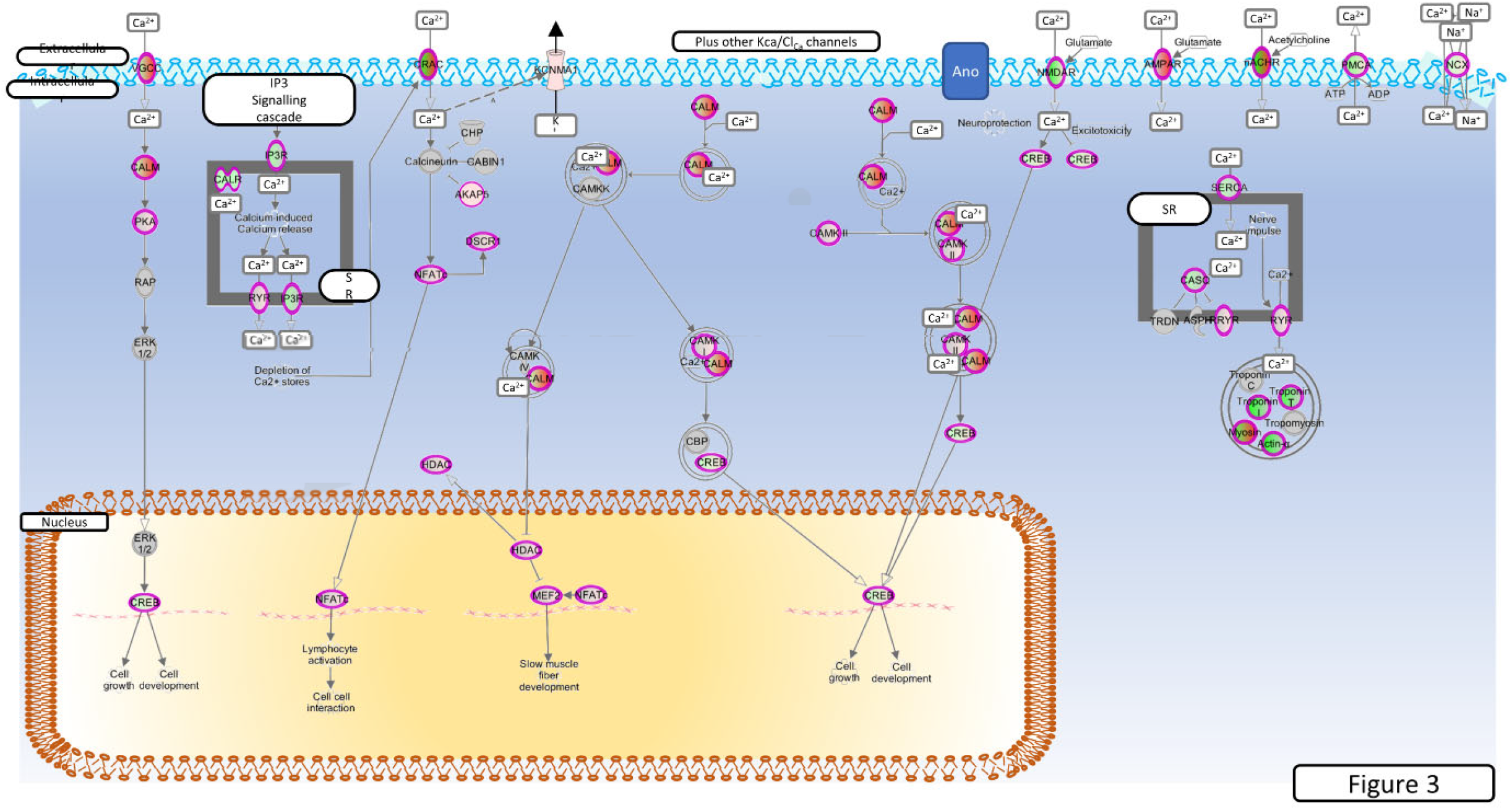
Enriched Calcium regulation pathway. Genes mapped to the calcium signalling canonical pathway by IPA. Red, increased expression; green, decreased expression (for interpretation of the references to colour in the figure legend, please refer to the online version of this article).

**Figure 4.**
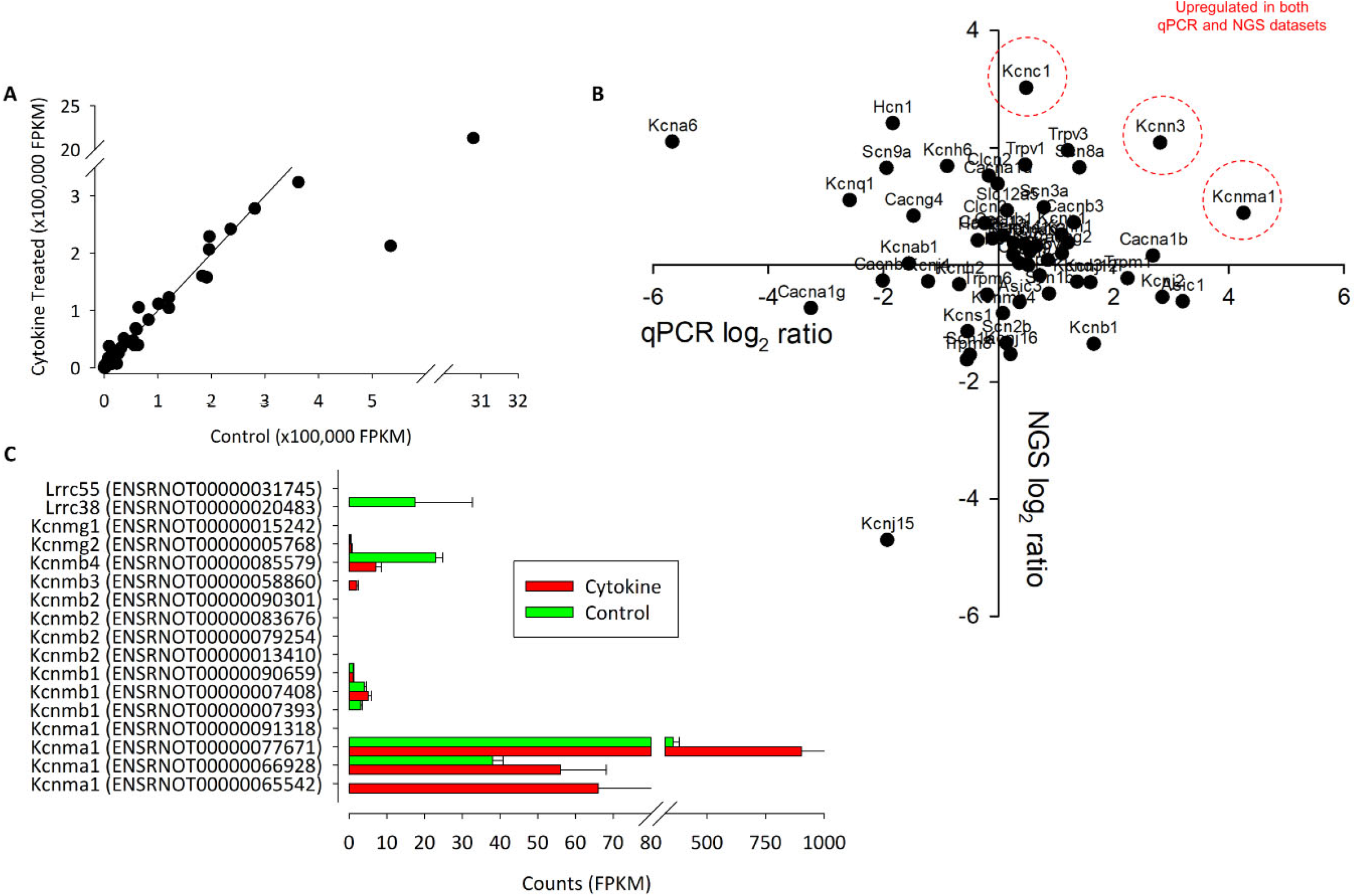
The correlation between Control and NGS expression data. **(A)** We found a strong correlation between control and treated (72 hrs of IL1β+TNFα) channel expression data. Each point represents the intersection of 4 control and 4 test values. In **(B),** the correlation between qPCR (ΔΔCts) and NGS ratio (FPKM values) are shown. Data suggest that there only a weak linear correlation between qPCR and NGS. P-value=0.3. R=0.29. n=14 (8 animals for NGS, 6 animals for PCR). Genes that were upregulated in both datasets are indicated in red. (C) Summary of NGS expression of all known (rat) splice variants of Kcnma1 and associated Kcnmb subunits.

### Next Generation RNA Sequencing Analysis of Ion Channel Gene Expression

Initial RNA-seq experiments were intended to give a transcriptome wide, unbiased, assessment of ion channel changes in cytokine treated FLS. We identified 190 channel genes, including porins, connexins and ion-channel isoform genes (including α- to ε-subunits), but excluded interacting proteins, other regulatory proteins and the so-called potassium tetramerization domain proteins. The top 50 genes, in terms of FPKM are given for control and cytokine treated datasets in Tables S3 and S4 respectively.

### Next Generation RNA Sequencing Ion Channel Differential Expression

One of the key ion channels involved with regulation of FLS is the large calcium activated potassium channel (BK, Kcnmx) and it is notable that there was a reduction in expression of β-subunit Kcnmb1/2 and appearance of Kcnmb3 after cytokine treatment. We further analysed this family transcript by transcript for the 13 known (already annotated) splice variants of these channels; Kcnma1, Kcnmb1, Kcnmb2, Kcnmb3, Kcnmb4 (Figure 4C). The most abundantly expressed of all these transcripts is Kcnma1 (ENSRNOT00000077671) with rather lower expression of any of the β-subunits and no detection of the one annotated Kcnmb4 variant. Following cytokine treatment expression of all of the detectable Kcnma1 splice variants was higher, but there was a lower abundance of Kcnmb4.

In total, 20 ion channel genes, undetectable in control conditions became detectable or ‘appeared’ after cytokine treatment (Table S5), of which, the top expressed of these was Trpc3. Conversely, 7 ion channel genes became undetectable or “disappeared” following cytokine treatment (Table S6). Following cytokine treatment, we found an additional 15 genes to be down by −1.5 (log_2_) or more and 21 genes 1.5 (log_2_) greater than control (Table 1 and Table 2 respectively). We used tissue from 4 animals for the NGS study each animal tissue split into test and control groups; the “*n*” presented in the legends refers to the number of biological replicates(= animals).

**Table 1:**
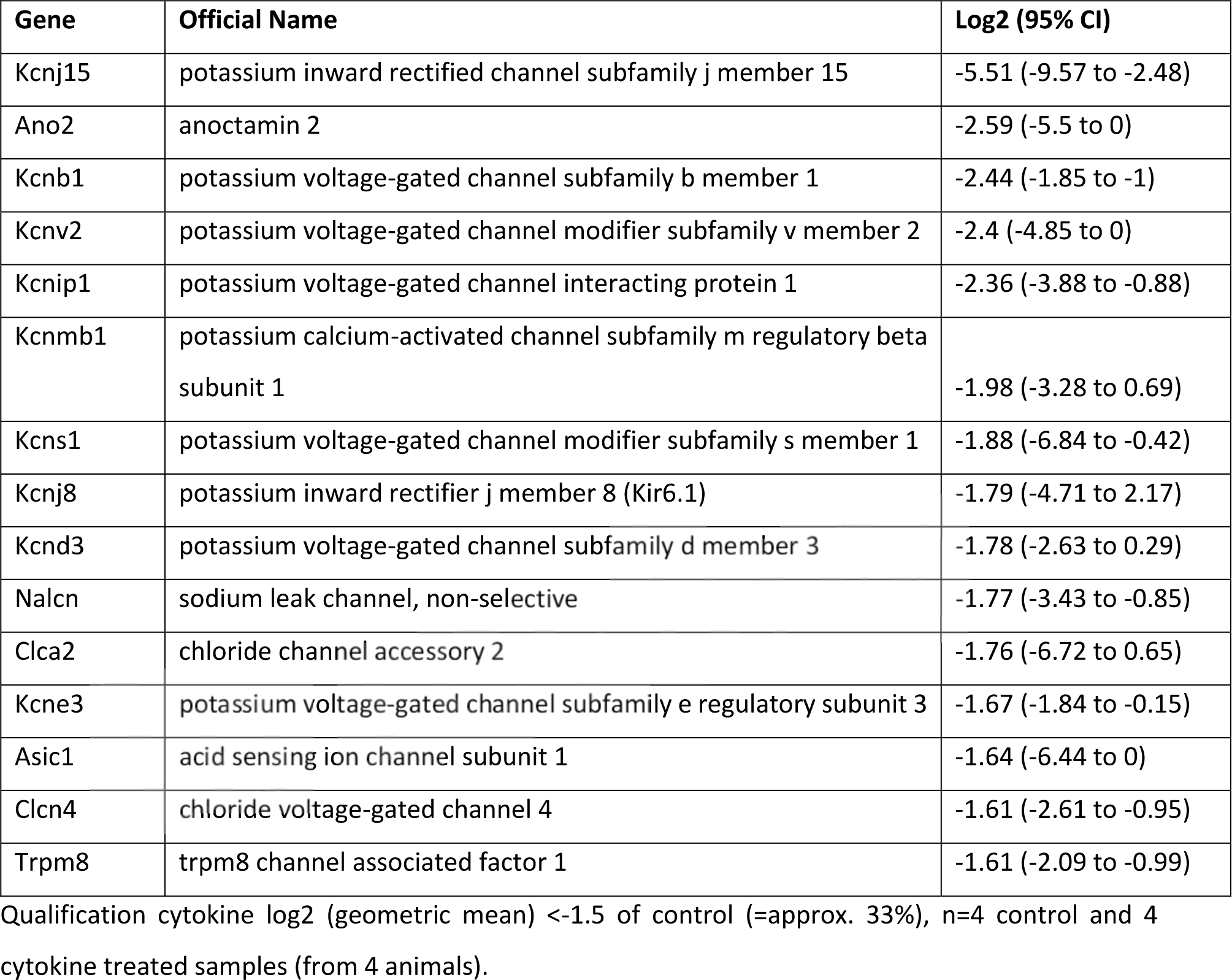
FLS channel gene RNA expression lower after cytokine (10ng/ml TNFα and IL1β) treatment.

**Table 2.** FLS channel gene RNA expression increased after cytokine (10ng/ml TNFα and IL1β) treatment.

| Gene ID | Official Name | Log2 (95% CI) |
| --- | --- | --- |
| Kcnc1 | potassium voltage-gated channel subfamily c member 1 | 4.32 (0.68 to 5.93) |
| Kcnc3 | potassium voltage-gated channel subfamily c member 3 | 3.46 (0.76 to 5.94) |
| Trpc3 | transient receptor potential cation channel subfamily c member 3 | 3.06 (3.73 to 6.5) |
| Catsper3 | cation channel sperm associated 3 | 2.83 (0.14 to 5.75) |
| Kcng3 | potassium voltage-gated channel modifier subfamily g member 3 | 2.81 (2.77 to 6.65) |
| Tmc7 | transmembrane channel like 7 | 2.77 (1.99 to 5.53) |
| Gjc3 | gap junction protein gamma 3 | 2.75 (0.03 to 4.35) |
| Scnn1g | sodium channel epithelial 1 gamma subunit | 2.56 (2.77 to 5.45) |
| Asic2 | acid sensing ion channel subunit 2 | 2.52 (0.66 to 4.65) |
| Kcnmb3 | potassium calcium-activated channel subfamily m regulatory beta subunit 3 | 2.34 (0 to 4.73) |
| Cracr2a | calcium release activated channel regulator 2a | 2.32 (0.01 to 1.96) |
| Kcnn3 | potassium calcium-activated channel subfamily n member 3 | 2.07 (0.57 to 4.09) |
| Tmc4 | transmembrane channel like 4 | 1.94 (0.27 to 2.47) |
| Asic4 | acid sensing ion channel subunit family member 4 | 1.75 (0.52 to 2.76) |
| Kcnj11 | potassium inward rectifier j member 11 (Kir6.2) | 1.74 (0 to 6.56) |
| Trpv6 | transient receptor potential cation channel subfamily v member 6 | 1.7 (0.01 to 5.43) |
| Hvcn1 | hydrogen voltage gated channel 1 | 1.69 (0.98 to 2.84) |
| Trpv1 | transient receptor potential cation channel subfamily v member 1 | 1.61 (0.98 to 2.07) |
| Kcnk7 | potassium two pore domain channel subfamily k member 7 | 1.6 (-1.85 to 1.03) |
| Cacna1d | calcium voltage-gated channel subunit alpha1 d | 1.57 (0.32 to 2.09) |
| Asic5 | acid sensing ion channel subunit family member 5 | 1.55 (0 to 5.78) |

Qualification cytokine log2 (geometric mean) >1.5 of control (=approx. 300%), n=4 control and 4 cytokine treated samples (from 4 animals).

### qPCR verification of RNA changes

To add further support to the unbiased RNA-seq ion channel analysis we performed qPCR on sets of control and IL1β/TNFα treated FLS with panels of Ca^2+^-potassium channels (Kcnma1, Kcnn1, Kcnn2, Kcnn3) and other ion channel genes (Figure 4B). We did not have primers for all the potassium channel genes identified by next generation sequencing. Three potassium genes were differentially expressed; two voltage-gated potassium channels Kcna6 and Kcnc2–significantly decreased (p<0.05, n=4,4), whereas the large calcium potassium channel Kcnma1 was upregulated (p<0.05, n=4,4).

### Electrophysiology

Neither RNA nor protein expression studies can confirm changes in functional ion channel expression or, functional changes resulting from post-translational changes, Furthermore, several different potassium channel isoforms were identified, of which some upregulated and some downregulated. Therefore, to investigate the effect of cytokine treatment on the electrophysiological fingerprint, we conducted functional assays of ion channel expression with patch-clamp electrophysiology. The primary changes observed in the more limited qRT-PCR data would predict over all *loss of* voltage-gated potassium ion channel activity and increase in the less voltage-dependent calcium activated potassium channels.

#### Resting membrane potential

Following cytokine treatment, the resting membrane potential of FLS decreased (depolarised) from −48.6±1.7 to −38.6±2.8mV, junction potential corrected, (data not shown, p<=0.05, unpaired t-test).

#### Current voltage currents

As seen in Figure 5A, some cells exhibited clear transient and sustained components whereas others exhibited only the sustained component of the current. We therefore analysed the transient and sustained components of the current separately in all cases.

We found no overall change in the maximum amplitude of the transient (Figure 5B), but sustained current (Figure 5C) density, measured at +20mV was decreased (12.5±2.3 pA/pF to 3.8±0.8pA/pF, n=24,20, p<0.05). There was no change in chord conductance measured at this potential (271±45 pS/pF 120±20 pS/pF, n=24,20). To characterise the nature of the conductance apparently *inhibited* by cytokine treatment we calculated the difference current for cytokine treatment (i.e., cytokine – treated current; Figure 5D). This revealed a strongly voltage-gated current with mid-point for activation 40±1.2 mV and slope 17.6±1.2 mV, *n*=25.

#### Pharmacological Modulation of currents

Despite the *increase* of KCNMA1 RNA in both qPCR and RNA-Seq our electrophysiological data show a reduced current density. To investigate if the maximum possible BK current is altered in cytokine treatment, we repeated our standard voltage protocol (above) in the presence of 1µM of the BK channel opener NS1619 and found a significant increase in current density in the presence of NS1619 in control cells (Figure 6A and 6C, p≤0.05, n=9,6), we did not see an equivalent increase in the cytokine treated cells (Figure 6B and 6D). We then repeated this experiment with 1µM of the BK channel inhibitor paxilline. Untreated cell current density was significantly smaller in the presence of paxilline, but not significantly reduced in treated cells (Figure 6E and F).

## Discussion

In this study we used an *in vitro* model of synovial cell inflammation to investigate the pathophysiology of cytokine induced synovitis. We have demonstrated that IL1β and TNFα treatment of FLS cells resulted in profound changes in arthritic, inflammatory and Ca^2+^ regulatory pathways similar to that reported in RA models. We found differential expression of several ion channels with transcriptomics and our electrophysiological experiments show a reduction of whole-cell current density.

### The pathophysiological validity of the 72hr IL1β and TNFα model

Treatment of joint tissue with TNFα and IL1β cytokines is an established in vitro model of inflammatory arthritis with tissue typically exposed to between 10ng/ml of TNFα and IL1β for between 2 to 7 days. In the present study, we use the lower end of the concentration range, 10ng and treat for 72hrs (De Ceuninck *et al.* 2004; Stevens *et al.* 2008; Pretzel *et al.* 2009; Stevens *et al.* 2009; Williams *et al.* 2011; Williams 2014). This regime is hypothesised to activate inflammatory pathways, but there will be no cytokine remaining by the time of the electrophysiological experiments that could cause confounding direct effects. We treated FLS cells with the pro-inflammatory cytokines IL1β and TNFα in order to understand the cellular changes that occur when these FLS are subjected to higher-than-normal levels of these cytokines *in vivo*, for example, in arthritis. This model has distinct advantages of 3Rs, consistency, reproducibility and allowing the investigation of distinct pathways in isolation but since it is an acute model it may lack some chronic features of *in vivo* models. Our transcriptome analysis demonstrated that pathways were activated in common with arthritis movement of cells, proliferation and both rheumatoid and osteoarthritis. In an *in vitro* model of RA, Tanner et al also showed up regulation of Kcnma1 (message and protein) (Tanner *et al.* 2019) analogous to that observed with our 72hr cytokine treatment. Furthermore, more recent data suggests a correlation between FLS invasiveness and expression of the β-subunit (KCNMB3) in human samples of FLS from RA patients (Petho *et al.* 2016). *Taken together with the changes in Ca^2+^ signalling, we show that our in vitro model captures several features of the inflammatory joint phenotype and that FLS have been “activated” as observed in animal models of arthritis*.

Causal analysis is a relatively new mathematical technique that allows one to move from probability of agreement or simple correlation towards probability of causation in networks. The IPA implementation of this identified both IL1β and TNFα as master-regulators of the changes we observe and considering our experimental design included time matched controls, this strongly supports a suitable dosage and incubation time. Whilst both IL1β and TNFα were identified by IPA as “master regulators”, TNFα transcriptome-wide causation was stronger than that of IL1β. Interestingly, FLS cells harvested from RA patients exhibit a marked transient elevation of intracellular Ca^2+^ on exposure to TNFα (Yoo *et al.* 2006) raising the possibility that this could be the initial trigger for the resulting pathway changes. However, it should be noted that such a transient lasts less than a minute and in our study cells were challenged with cytokines 72hrs prior to experiments. Furthermore, cells were replaced in cytokine-free medium for electrophysiology, so there would be no cytokine physically present at that time. Also, the previously reported TNFα induced Ca^2+^ signal was only clear in FLS from RA; it was largely absent in FLS from OA patients and not investigated in FLS from healthy controls (Yoo *et al.* 2006).

### Previous studies of differential expression of membrane ion channels in synovium

Until very recently, little was known of the FLS ion channel compliment (the “channelome”) compared to that of the another central joint cell, the chondrocyte (Barrett-Jolley *et al.* 2010). One of the best-studied families of ion channels in FLS, however, is the Ca^2+^-activated potassium channel family. The high conductance member of this family, termed BK (KCa1.1 or KCNMA1) and the “intermediate” conductance member (“IK”, or KCa3.1) are both expressed and have roles in invasive migration, proliferation, cytokine and MMP release (Hu *et al.* 2012; Friebel *et al.* 2014). Interestingly, inhibitors of BK decrease the signs of joint degeneration in the pristane-induced arthritis model. These channels are therefore potential drug targets to protect against joint degeneration as well as being putative biomarkers. Whilst the BK channel β-subunit (KCNMB1) was slightly increased in transcript abundance in the Lambert *et al.,* 2014 data (similar seen by Huber et al., 2008), both of the two BK α-subunit (KNMA1) probes on the chip exhibit small decreases in expression. It should be noted that the expression of BK channel β-subunits confers modulation of ion channel activity, in many cases decreasing its sensitivity to, for example, Ca^2+^ ions (Lippiat *et al.* 2003; Mobasheri *et al.* 2012).

A recent study by Kondo et al demonstrated that human FLS express high levels of both intermediate (K_Ca_3.1) and large (BK/K_Ca_1.1/KCNMA1) Ca^2+^-activated potassium channels. The other most highly expressed ion channels identified by Kondo et al (2018) were KCNK2, ANO6, ANO10 and KCNK6. KCNK2/6 are members of the two-pore-domain potassium channel family and are particularly thought of as molecular sensors, whereas the ANO (anoctamin) channels are members of the large chloride channel family. This family is relatively understudied compared to potassium channels, but ANO6 (TMEM16F) is, interestingly, thought to be a Ca^2+^-activated chloride channel as well as a lipid “scramblase” (Scudieri *et al.* 2015), therefore, likely to be activated in parallel to Ca^2+^-activated potassium channels.

### Changes in ion channel currents in the present study

The phenotype of FLS current recorded by whole-cell patch clamp was quite variable in terms of the presence of transient and sustained components of current, as seen in Figure 5. We found no significant changes in the transient phase of current and therefore, we focussed on the sustained component of voltage activated currents that would be expected to include BK activity if it was present. Although we found a significant increase in RNA expression of the BK α-subunit gene, Kcnma1, following cytokine treatment (by qPCR), there was a decrease in over-all current density of the sustained current. To characterise the voltage-gated characteristics of the current *lost* after cytokine treatment we subtracted IV curves following cytokine treatment from the control IV curves and transformed this to a conductance-voltage curve. Since this curve fully saturated, we were able to fit it with a Boltzmann curve and derive the midpoint for activation; +40mV. This is rather positive to most voltage-gated potassium channels, the most “positive” of which (Kcnbx/Kcncx) have activation mid-points in the +10 to +20mV range (Coetzee *et al.* 1999). This cytokine sensitive conductance could be an ensemble average of a number of different ion channel changes, including increase of some and decrease of others. It could also be a BK current under conditions of low intracellular Ca^2+^, (Cui *et al.* 1997; Coetzee *et al.* 1999; Contreras *et al.* 2012)(Cui *et al.* 2009) but, since the true local concentration of intracellular Ca^2+^ is unknown, this is difficult to assess. V_1/2_ for the BK channel in the virtual absence of Ca^2+^ can be much higher, for example 200-300mV (Bao and Cox 2005; Orio and Latorre 2005; Wang and Brenner 2006). The value of this parameter also depends on the nature of the co-expressed β-subunit (Coetzee *et al.* 1999; Contreras *et al.* 2012). V_1/2_ is typically shifted to the left in the presence of beta subunits (Brenner 2014). In our transcript data (Figure 4c) we show a range of BK transcripts including the Lrrcx subunits, but none, on their own, show significant alteration by treatment. Note that with experimentally elevated intracellular Ca^2+^ concentration these would likely lie well to the left of where they lay in our experiments. In our experiments we included 5mM EGTA, which allows for the Ca^2+^-activation of BK channels so long as they are physically close to the Ca^2+^ source (Fakler and Adelman 2008). Functional coupling between BK and the Ca^2+^ source appears a common phenomenon (Fakler and Adelman 2008); we also showed this with a coupling between TRPV4 channels and KCa channels previously in neurones (Feetham *et al.* 2015) and it has also been shown in smooth muscle cells too (Nilius and Droogmans 2001). Increasing, or attempting to “clamp” intracellular Ca^2+^ before investigating BK channels would be tempting, but this could cause greater constitutive activation of BK and mask a physiological coupling.

**Figure 5.**
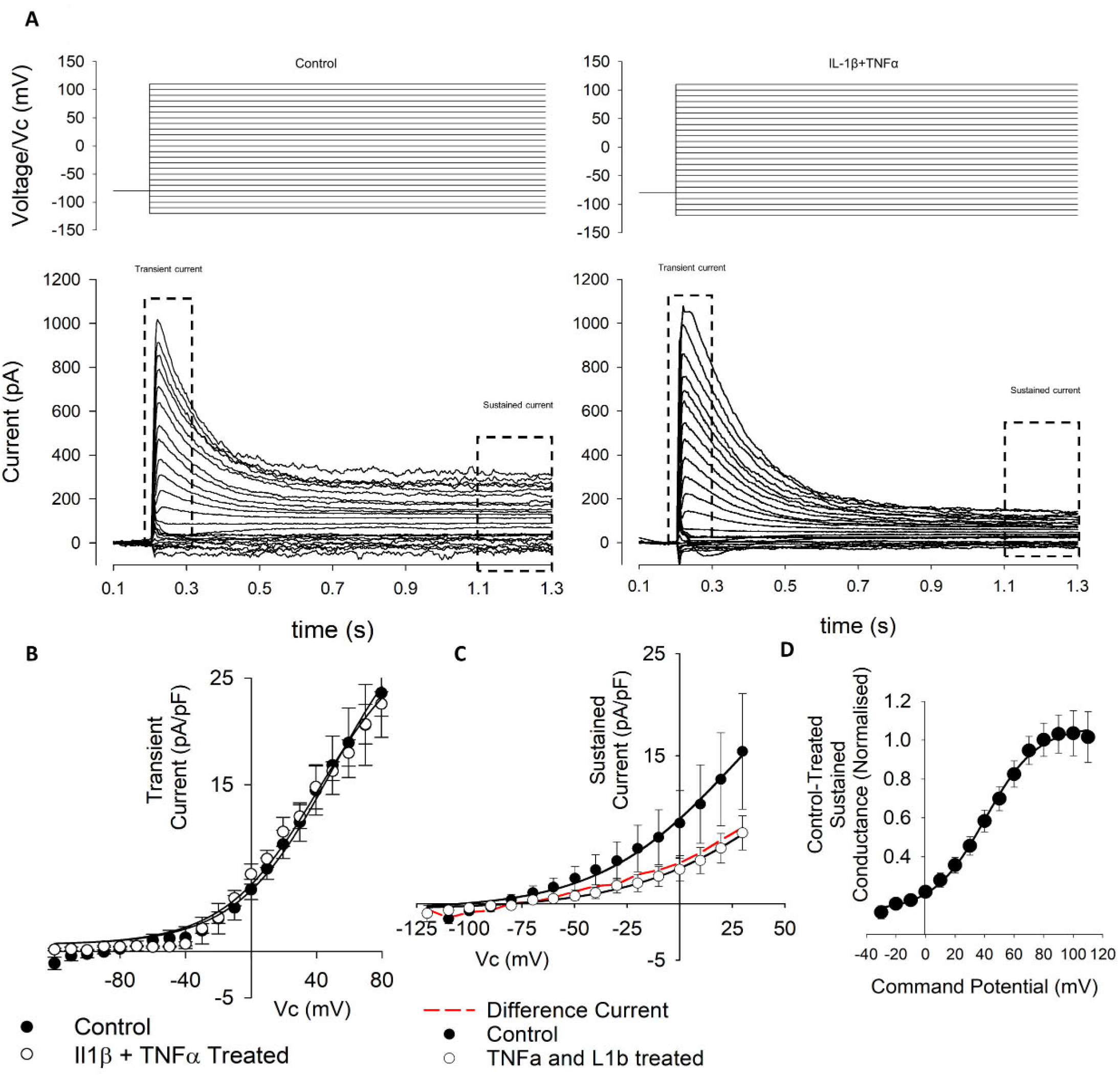
Whole-cell voltage-gated currents from control and cytokine treated FLS. **(A)** Top panels show the voltage step protocols and the evoked currents are shown below for control (left) and IL1β+TNFα (right) conditions. Note that FLS exhibit both transient and sustained currents. These phases of current were then analysed separately as indicated. **(B)** Current-voltage curves (left) from the *transient* currents recorded in a number or experiments such as that illustrated in (A). There was no significant difference between control and treated *transient* current density. Data points are shown as mean± SEM (n=18 for control and n=11 for IL1β+TNFα). (C) Current-voltage curves from the *sustained* currents recorded in a number or experiments such as that illustrated in (A). The red line in the current-voltage curve is the difference current for control-cytokine treated. **(D)** Difference conductance-voltage curve for the cytokine difference current shown in (C). The line is fit with a Boltzmann, *see text*.

**Figure 6.**
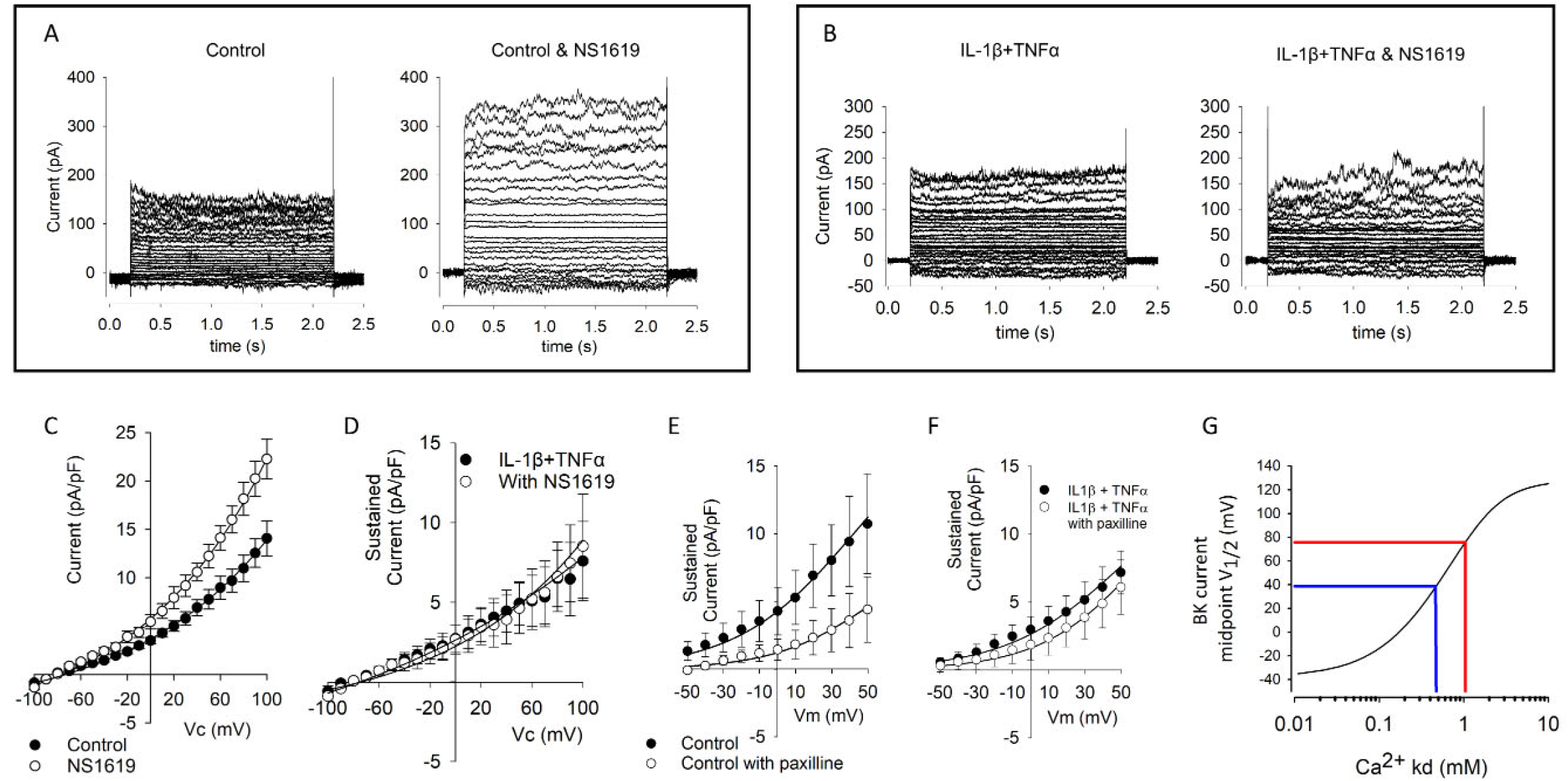
Effects of BK channel drugs on FLS whole-cell currents. **(A)** Representative examples of untreated (control) cells before (left) and in the presence of 1µM of the BK channel opener NS1619 (left). Voltage protocols as shown in Figure 5. **(B)** Representative raw example current families of cytokine (10ng/ml IL1β+TNFα) treated cells in the absence (left) and presence (right) of 1µM NS1619. **(C)** Current-voltage curves from a number of control FLS cells such as that shown in (A). Current is significantly greater in the presence of NS1619, p<0.05, *n=* 9 and 6. **(D)** Current-voltage curves for a number of treated FLS cells such as that shown in (B). These two curves are not significantly different from each other *n=* 6 and 6. **(E)** Current-voltage curves from a number of control FLS cells treated with and without paxilline; there is a significant decrease in paxilline current p<0.05 n=13,5), but **(F)** there was no significant different of current density in the presence of the BK inhibitor paxilline following cytokine treatment (n=6, 12). **(G)** *Numerical simulation of data from Clark et al 2017 model.* To quantify the degree of change of BK channel modulation apparently taking place with this treatment we used the model of verbatim with the exception that we varied the inherent BK channel Ca^2+^ sensitivity parameter *kd.* In this simulation, the independent variable *kd* is plotted on the x-axis and the predicted BK current midpoint (V_1/2_) is plotted on the y-axis. In blue we have added hypothetical midpoints of 40mV and 80mV representing a hypothetical shift in 40mV by cytokine treatment. The complete MATLAB code for this simulation is included in the supplementary data (2).

Our electrophysiological experiments also demonstrated significantly more depolarised resting membrane potentials in cytokine treated cells. This could result from loss of constitutive potassium or chloride conductances, but equally it could result from the elevation of the nonspecific or Na^+^ selective or ion channels such as Trpc3 or Asic2 etc. (Table II).

The BK pharmacological activator (NS1619) and inhibitor (paxilline) both had the expected effects (increase and decrease of current density respectively) in the untreated FLS but neither had a significant effect. This is surprising in the light of the Tanner et al data showing that paxilline ameliorate development joint degeneration in a model of RA (Tanner *et al.* 2015) and the continued effectiveness of paxilline in FLS from RA patients (Petho *et al.* 2016). One possible explanation could be differences in transcript expression between strains as has been observed with ion channels previously (Kunert-Keil *et al.* 2006). Our RNA data showed a clear shift from RNA-expression of β1 (and β2) subunits to β3 and sensitivity to voltage, calcium and some drugs is well-known to be conveyed by co-expression of the β subunits (McManus *et al.* 1995; Uebele *et al.* 2000; Yang *et al.* 2009). However, neither paxilline nor NS1619 themselves are thought to be influenced by the β-subunit; paxilline acts as a closed channel blocker (Zhou and Lingle 2014) whereas NS1619 opens the BK channel by binding to the KCNMA1 S6/RCK linker and is effective in some splice variants, but not others (Soom *et al.* 2008; Gessner *et al.* 2012). In humans there are 93 known s[lice variants of KCNMA1, but there are only 4 in rats listed on ENSEMBL (Zerbino *et al.* 2018). Figure 4 shows the relative transcript expression for Kcnma1. The most abundant transcript (ENSRNOT00000077671 Kcnma1-203) is present in both conditions. None of the Kcnma1 transcripts decrease with cytokine so it seems unlikely (but not impossible) that pharmacological changes result directly from splice variant switching. Note that the one known transcript that was not detected is the truncated transcript Kcnma1-204 (ENSRNOT00000091318). The lack of effect of NS1619 and paxilline after cytokine treatment could be also be secondary to changes in β-subunits or intracellular Ca^2+^ in microdomains or some unknown reason leaving insufficient residual functional BK activity to be noticeably modulated. The simple (K^+^ channel focussed) FLS electrophysiological model of Clark allows us to use numerical simulation to estimate the change in BK channel Ca^2+^ that would be required to shift a BK conductance voltage curve by about +40mV and largely leave the cells virtually free of measurable BK current. We find that retaining all the parameters of (Clark *et al.* 2017) except the Ca^2+^ *Kd* itself, such data would be the equivalent to increasing *Kd* from 0.46µM to approximately 1.05µM. The upstream pathway (beyond the activation of the cytokine pathway including NFκB etc) for these changes is difficult to identify from our data, especially since there are relatively few ion channel interaction data in the IPA databases. It is notable however, that there were significant changes in the calcium signalling pathway (Figure 3) and so it is possible that change in calcium activated potassium channel expression follows this, by way of compensatory expression. The lack of apparent effect of paxilline could also be technical. For example, there was considerable variability between cells and we compared population means rather than pairing each cell with and without paxilline increasing the risk of a type II error.

### Role of ion channels in pro-inflammatory cytokine production and secretion

Ion channels are involved not just in the response to cytokines, but can also contribute to their production. For example, ionotropic NMDA and kainite glutamate receptors contribute to synovial inflammation by increasing expression of the inflammatory cytokine IL-6 (Flood *et al.* 2007) and nicotinic acetylcholine receptor activation reduces the synovial production of IL-6, IL-8, TNFα and several other cytokines (Waldburger *et al.* 2008; van Maanen *et al.* 2009). P2X7 is an established conduit for release of mediators such as IL1β (Mortaz *et al.* 2012). We saw little P2X7 RNA, but there is evidence that in chondrocytes P2X1, may sub serve the same function (Varani *et al.* 2008) and we detected RNA for this channel only after cytokine treatment. In a previous report, inhibition of the small Ca^2+^ activated potassium ion channel decreased the production of cytokines IL-6, IL-8 and MCP1 in response to TGF-1β, but they did not examine secretion of IL1β or TNFα (Friebel *et al.* 2014). In other-words activation of Ca^2+^ potassium ion channels is essentially a secretion trigger. This result is somewhat counter intuitive since one would expect activation of a potassium channel to hyperpolarise and decrease secretion. One possible explanation is that activation of Ca^2+^ potassium ion channels draws in additional Ca^2+^ by increasing the driving force for Ca^2+^ entry.

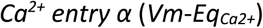

Where negative values are equivalent to inward driving force for Ca^2+^, *Vm* is the membrane potential and *Eq*_*Ca2+*_ is the equilibrium potential for Ca^2+^.

## Conclusion

To our knowledge, this is the first report using a combined NGS and patch-clamp electrophysiological approach to understanding the control of potassium channels in inflammation in joint tissues. We found an increased RNA expression of the BK potassium gene Kcnma1, but the constitutive activity of the channel was not increased. The decreased sensitivity to voltage activation and to drugs could be explained by a switch the RNA expression of β-subunits KcnmB1, 2 and 3.

## Acknowledgements

For funding, the authors would like to thank the European Union’s Seventh Framework Programme for research, technological development and demonstration (grant agreement No. 305815) and King Abdulaziz University, Jeddah, Saudi Arabia.

## Author Contributions

- Conception and design All authors.
- Analysis and interpretation of the data All authors
- Drafting of the article RBJ/FMOB/KK
- Critical revision of the article for important intellectual content All authors
- Final approval of the article All authors
- Obtaining of funding RBJ AM OH

## References

Afgan, E., Baker, D., Batut, B., van den Beek, M., Bouvier, D., Cech, M., Chilton, J., Clements, D., Coraor, N., Gruning, B.A., Guerler, A., Hillman-Jackson, J., Hiltemann, S., Jalili, V., Rasche, H., Soranzo, N., Goecks, J., Taylor, J., Nekrutenko, A. and Blankenberg, D. (2018) ‘The Galaxy platform for accessible, reproducible and collaborative biomedical analyses: 2018 update’, Nucleic Acids Res, 46(W1), W537–W544, available: http://dx.doi.org/10.1093/nar/gky379.

Bao, L. and Cox, D.H. (2005) ‘Gating and ionic currents reveal how the BKCa channel’s Ca2+ sensitivity is enhanced by its beta1 subunit’, J Gen Physiol, 126(4), 393–412, available: http://dx.doi.org/10.1085/jgp.200509346.

Barrett-Jolley, R., Lewis, R., Fallman, R. and Mobasheri, A. (2010) ‘The Emerging Chondrocyte Channelome’, Front Physiol, 1(135), 1–11, available: http://dx.doi.org/10.3389/fphys.2010.00135.

Bartok, B. and Firestein, G.S. (2010) ‘Fibroblast-like synoviocytes: key effector cells in rheumatoid arthritis’, Immunol Rev, 233(1), 233–55, available: http://dx.doi.org/10.1111/j.0105-2896.2009.00859.x.

Berenbaum, F. (2013) ‘Osteoarthritis as an inflammatory disease (osteoarthritis is not osteoarthrosis!)’. Osteoarthritis Cartilage, 21(1), 16–21, available: http://dx.doi.org/10.1016/j.joca.2012.11.012.

Brenner, R. (2014) ‘Knockout of the BK β2 subunit reveals the importance of accessorizing your channel’, J Gen Physiol, 144(5), 351–6, available: http://dx.doi.org/10.1085/jgp.201411291.

Clark, R.B., Schmidt, T.A., Sachse, F.B., Boyle, D., Firestein, G.S. and Giles, W.R. (2017) ‘Cellular electrophysiological principles that modulate secretion from synovial fibroblasts’, J Physiol, 595(3), 635–645, available: http://dx.doi.org/10.1113/JP270209.

Coetzee, W.A., Amarillo, Y., Chiu, J., Chow, A., Lau, D., McCormack, T., Moreno, H., Nadal, M.S., Ozaita, A., Pountney, D., Saganich, M., Vega-Saenz de Miera, E. and Rudy, B. (1999) ‘Molecular diversity of K+ channels’, Ann NY Acad Sci, 868, 233–85.

Contreras, G.F., Neely, A., Alvarez, O., Gonzalez, C. and Latorre, R. (2012) ‘Modulation of BK channel voltage gating by different auxiliary beta subunits’, Proceedings of the National Academy of Sciences of the United States of America, 109(46), 18991–6.

Cui, J., Cox, D.H. and Aldrich, R.W. (1997) ‘Intrinsic voltage dependence and Ca2+ regulation of mslo large conductance Ca-activated K+ channels’, J Gen Physiol, 109(5), 647–73.

Cui, J., Yang, H. and Lee, U.S. (2009) ‘Molecular mechanisms of BK channel activation’, Cell Mol Life Sci, 66(5), 852–75, available: http://dx.doi.org/10.1007/s00018-008-8609-x.

De Ceuninck, F., Dassencourt, L. and Anract, P. (2004) ‘The inflammatory side of human chondrocytes unveiled by antibody microarrays’, Biochem Biophys Res Commun, 323(3), 960–9, available: http://dx.doi.org/10.1016/j.bbrc.2004.08.184.

Fakler, B. and Adelman, J.P. (2008) ‘Control of K(Ca) channels by calcium nano/microdomains’, Neuron, 59(6), 873–81, available: http://dx.doi.org/10.1016/j.neuron.2008.09.001.

Feetham, C.H., Nunn, N., Lewis, R., Dart, C. and Barrett-Jolley, R. (2015) ‘TRPV4 and KCa ion channels functionally couple as osmosensors in the paraventricular nucleus’, Br J Pharmacol, 172(7), 1753–68, available: http://dx.doi.org/10.1111/bph.13023.

Fernandez-Madrid, F., Karvonen, R.L., Teitge, R.A., Miller, P.R., An, T. and Negendank, W.G. (1995) ‘Synovial thickening detected by MR imaging in osteoarthritis of the knee confirmed by biopsy as synovitis’, Magnetic Resonance Imaging, 13(2), 177–183, available: http://dx.doi.org/https://doi.org/10.1016/0730-725X(94)00119-N.

Flood, S., Parri, R., Williams, A., Duance, V. and Mason, D. (2007) ‘Modulation of interleukin-6 and matrix metalloproteinase 2 expression in human fibroblast-like synoviocytes by functional ionotropic glutamate receptors’, Arthritis Rheum, 56(8), 2523–34, available: http://dx.doi.org/10.1002/art.22829.

Friebel, K., Schonherr, R., Kinne, R.W. and Kunisch, E. (2014) ‘Functional role of the KCa3.1 potassium channel in synovial fibroblasts from rheumatoid arthritis patients’, J Cell Physiol, available: http://dx.doi.org/10.1002/jcp.24924.

Gessner, G., Cui, Y.M., Otani, Y., Ohwada, T., Soom, M., Hoshi, T. and Heinemann, S.H. (2012) ‘Molecular mechanism of pharmacological activation of BK channels’, Proc Natl Acad Sci USA, 109(9), 3552–7, available: http://dx.doi.org/10.1073/pnas.1114321109.

Grubb, B.D. (2004) ‘Activation of sensory neurons in the arthritic joint’, Novartis Found Symp, 260, 28–36; discussion 36-48, 100-4, 277-9.

Hu, X., Laragione, T., Sun, L., Koshy, S., Jones, K.R., Ismailov, II, Yotnda, P., Horrigan, F.T., Gulko, P.S. and Beeton, C. (2012) ‘KCa1.1 potassium channels regulate key proinflammatory and invasive properties of fibroblast-like synoviocytes in rheumatoid arthritis’, Journal of Biological Chemistry, 287(6), 4014–22, available: http://dx.doi.org/10.1074/jbc.M111.312264.

Kahlenberg, J.M. and Fox, D.A. (2011) ‘Advances in the medical treatment of rheumatoid arthritis’, Hand Clin, 27(1), 11–20, available: http://dx.doi.org/10.1016/j.hcl.2010.09.002.

Kumahashi, N., Naitou, K., Nishi, H., Oae, K., Watanabe, Y., Kuwata, S., Ochi, M., Ikeda, M. and Uchio, Y. (2011) ‘Correlation of changes in pain intensity with synovial fluid adenosine triphosphate levels after treatment of patients with osteoarthritis of the knee with high-molecular-weight hyaluronic acid’, Knee, 18(3), 160–4, available: http://dx.doi.org/10.1016/j.knee.2010.04.013.

Kunert-Keil, C., Bisping, F., Kruger, J. and Brinkmeier, H. (2006) ‘Tissue-specific expression of TRP channel genes in the mouse and its variation in three different mouse strains’, Bmc Genomics, 7, 159, available: http://dx.doi.org/10.1186/1471-2164-7-159.

Large, R.J., Hollywood, M.A., Sergeant, G.P., Thornbury, K.D., Bourke, S., Levick, J.R. and McHale, N.G. (2010) ‘Ionic currents in intimal cultured synoviocytes from the rabbit’, Am J Physiol Cell Physiol, 299(5), C1180–94, available: http://dx.doi.org/10.1152/ajpcell.00028.2010.

Lewis, R., Feetham, C.H., Gentles, L., Penny, J., Tregilgas, L., Tohami, W., Mobasheri, A. and Barrett-Jolley, R. (2013) ‘Benzamil sensitive ion channels contribute to volume regulation in canine chondrocytes’, British Journal of Pharmacology, 168(7), 1584–1596, available: http://dx.doi.org/10.1111/j.1476-5381.2012.02185.x.

Lippiat, J.D., Standen, N.B., Harrow, I.D., Phillips, S.C. and Davies, N.W. (2003) ‘Properties of BK(Ca) channels formed by bicistronic expression of hSloalpha and beta1-4 subunits in HEK293 cells’, The Journal of membrane biology, 192(2), 141–8.

Martel-Pelletier, J., Alaaeddine, N. and Pelletier, J.P. (1999) ‘Cytokines and their role in the pathophysiology of osteoarthritis’, Front Biosci, 4, D694–703.

Mathiessen, A. and Conaghan, P.G. (2017) ‘Synovitis in osteoarthritis: current understanding with therapeutic implications’, Arthritis Res Ther, 19(1), 18, available: http://dx.doi.org/10.1186/s13075-017-1229-9.

McManus, O.B., Helms, L.M., Pallanck, L., Ganetzky, B., Swanson, R. and Leonard, R.J. (1995) ‘Functional role of the beta subunit of high conductance calcium-activated potassium channels’, Neuron, 14(3), 645–50.

Mobasheri, A., Gent, T.C., Womack, M.D., Carter, S.D., Clegg, P.D. and Barrett-Jolley, R. (2005) ‘Quantitative analysis of voltage-gated potassium currents from primary equine (Equus caballus) and elephant (Loxodonta africana) articular chondrocytes’, American Journal of Physiology-Regulatory Integrative and Comparative Physiology, 289(1), R172–R180, available: http://dx.doi.org/10.1152/ajpregu.00710.2004.

Mobasheri, A., Lewis, R., Ferreira-Mendes, A., Rufino, A., Dart, C. and Barrett-Jolley, R. (2012) ‘Potassium channels in articular chondrocytes’, Channels, 6(6), 416–425, available: http://dx.doi.org/http://dx.doi.org/10.4161/chan.22340.

Mortaz, E., Adcock, I.M., Shafei, H., Masjedi, M.R. and Folkerts, G. (2012) ‘Role of P2X7 Receptors in Release of IL-1β: A Possible Mediator of Pulmonary Inflammation’, Tanaffos, 11(2), 6–11.

Nilius, B. and Droogmans, G. (2001) ‘Ion channels and their functional role in vascular endothelium’, Physiological Reviews, 81(4), 1415–1459.

Noss, E.H. and Brenner, M.B. (2008) ‘The role and therapeutic implications of fibroblast-like synoviocytes in inflammation and cartilage erosion in rheumatoid arthritis’, Immunol Rev, 223, 252–70, available: http://dx.doi.org/10.1111/j.1600-065X.2008.00648.x.

Orio, P. and Latorre, R. (2005) ‘Differential effects of beta 1 and beta 2 subunits on BK channel activity’, J Gen Physiol, 125(4), 395–411, available: http://dx.doi.org/10.1085/jgp.200409236.

Petho, Z., Tanner, M.R., Tajhya, R.B., Huq, R., Laragione, T., Panyi, G., Gulko, P.S. and Beeton, C. (2016) ‘Different expression of beta subunits of the KCa1.1 channel by invasive and non-invasive human fibroblast-like synoviocytes’, Arthritis Res Ther, 18(1), 103, available: http://dx.doi.org/10.1186/s13075-016-1003-4.

Pretzel, D., Pohlers, D., Weinert, S. and Kinne, R.W. (2009) ‘In vitro model for the analysis of synovial fibroblast-mediated degradation of intact cartilage’, Arthritis Res Ther, 11(1), R25, available: http://dx.doi.org/10.1186/ar2618.

Roemer, F.W., Felson, D.T., Yang, T., Niu, J., Crema, M.D., Englund, M., Nevitt, M.C., Zhang, Y., Lynch, J.A., El Khoury, G.Y., Torner, J., Lewis, C.E. and Guermazi, A. (2013) ‘The association between meniscal damage of the posterior horns and localized posterior synovitis detected on T1-weighted contrast-enhanced MRI—The MOST study’, Seminars in Arthritis and Rheumatism, 42(6), 573–581, available: http://dx.doi.org/https://doi.org/10.1016/j.semarthrit.2012.10.005.

Scudieri, P., Caci, E., Venturini, A., Sondo, E., Pianigiani, G., Marchetti, C., Ravazzolo, R., Pagani, F. and Galietta, L.J.V. (2015) ‘Ion channel and lipid scramblase activity associated with expression of TMEM16F/ANO6 isoforms’, Journal of Physiology-London, 593(17), 3829–3848, available: http://dx.doi.org/10.1113/Jp270691.

Sellam, J. and Berenbaum, F. (2010) ‘The role of synovitis in pathophysiology and clinical symptoms of osteoarthritis’, Nat Rev Rheumatol, 6(11), 625–35, available: http://dx.doi.org/10.1038/nrrheum.2010.159.

Soom, M., Gessner, G., Heuer, H., Hoshi, T. and Heinemann, S.H. (2008) ‘A mutually exclusive alternative exon of slo1 codes for a neuronal BK channel with altered function’, Channels (Austin), 2(4), 278–82, available: http://dx.doi.org/10.4161/chan.2.4.6571.

Stevens, A.L., Wishnok, J.S., Chai, D.H., Grodzinsky, A.J. and Tannenbaum, S.R. (2008) ‘A sodium dodecyl sulfate-polyacrylamide gel electrophoresis-liquid chromatography tandem mass spectrometry analysis of bovine cartilage tissue response to mechanical compression injury and the inflammatory cytokines tumor necrosis factor alpha and interleukin-1beta’, Arthritis Rheum, 58(2), 489–500, available: http://dx.doi.org/10.1002/art.23120.

Stevens, A.L., Wishnok, J.S., White, F.M., Grodzinsky, A.J. and Tannenbaum, S.R. (2009) ‘Mechanical injury and cytokines cause loss of cartilage integrity and upregulate proteins associated with catabolism, immunity, inflammation, and repair’, Mol Cell Proteomics, 8(7), 1475–89, available: http://dx.doi.org/10.1074/mcp.M800181-MCP200.

Sutton, S., Clutterbuck, A., Harris, P., Gent, T., Freeman, S., Foster, N., Barrett-Jolley, R. and Mobasheri, A. (2009) ‘The contribution of the synovium, synovial derived inflammatory cytokines and neuropeptides to the pathogenesis of osteoarthritis’, Veterinary Journal, 179(1), 10–24, available: http://dx.doi.org/10.1016/j.tvjl.2007.08.013.

Tanner, M.R., Hu, X., Huq, R., Tajhya, R.B., Sun, L., Khan, F.S., Laragione, T., Horrigan, F.T., Gulko, P.S. and Beeton, C. (2015) ‘KCa1.1 Inhibition Attenuates Fibroblast-like Synoviocyte Invasiveness and Ameliorates Disease in Rat Models of Rheumatoid Arthritis’, Arthritis Rheumatol, 67(1), 96–106, available: http://dx.doi.org/10.1002/art.38883.

Tanner, M.R., Pennington, M.W., Chauhan, S.S., Laragione, T., Gulko, P.S. and Beeton, C. (2019) ‘KCa1.1 and Kv1.3 channels regulate the interactions between fibroblast-like synoviocytes and T lymphocytes during rheumatoid arthritis’, Arthritis Res Ther, 21(1), 6, available: http://dx.doi.org/10.1186/s13075-018-1783-9.

Uebele, V.N., Lagrutta, A., Wade, T., Figueroa, D.J., Liu, Y., McKenna, E., Austin, C.P., Bennett, P.B. and Swanson, R. (2000) ‘Cloning and functional expression of two families of beta-subunits of the large conductance calcium-activated K+ channel’, J Biol Chem, 275(30), 23211–8, available: http://dx.doi.org/10.1074/jbc.M910187199.

van Maanen, M.A., Stoof, S.P., van der Zanden, E.P., de Jonge, W.J., Janssen, R.A., Fischer, D.F., Vandeghinste, N., Brys, R., Vervoordeldonk, M.J. and Tak, P.P. (2009) ‘The alpha7 nicotinic acetylcholine receptor on fibroblast-like synoviocytes and in synovial tissue from rheumatoid arthritis patients: a possible role for a key neurotransmitter in synovial inflammation’, Arthritis Rheum, 60(5), 1272–81, available: http://dx.doi.org/10.1002/art.24470.

Varani, K., De Mattei, M., Vincenzi, F., Tosi, A., Gessi, S., Merighi, S., Pellati, A., Masieri, F., Ongaro, A. and Borea, P.A. (2008) ‘Pharmacological characterization of P2X1 and P2X3 purinergic receptors in bovine chondrocytes’, Osteoarthritis Cartilage, 16(11), 1421–9, available: http://dx.doi.org/10.1016/j.joca.2008.03.016.

Waldburger, J.M., Boyle, D.L., Pavlov, V.A., Tracey, K.J. and Firestein, G.S. (2008) ‘Acetylcholine regulation of synoviocyte cytokine expression by the alpha7 nicotinic receptor’, Arthritis Rheum, 58(11), 3439–49, available: http://dx.doi.org/10.1002/art.23987.

Wang, B. and Brenner, R. (2006) ‘An S6 mutation in BK channels reveals beta1 subunit effects on intrinsic and voltage-dependent gating’, J Gen Physiol, 128(6), 731–44, available: http://dx.doi.org/10.1085/jgp.200609596.

Williams, A. (2014) ‘Proteomic studies of an explant model of equine articular cartilage in response to proinflammatory and anti-inflammatory stimuli’, PhD Thesis University of Nottingham.

Williams, A., Smith, J.R., Allaway, D., Harris, P., Liddell, S. and Mobasheri, A. (2011) ‘Strategies for optimising proteomic studies of the cartilage secretome: establishing the time course for protein release and evaluating responses of explant cultures to il-1 beta, tnf-alpha and carprofen’, Osteoarthritis and Cartilage, 19, S209–S209.

Yang, C.T., Zeng, X.H., Xia, X.M. and Lingle, C.J. (2009) ‘Interactions between beta subunits of the KCNMB family and Slo3: beta4 selectively modulates Slo3 expression and function’. PLoS ONE, 4(7), e6135, available: http://dx.doi.org/10.1371/journal.pone.0006135.

Yoo, S.A., Park, B.H., Park, G.S., Koh, H.S., Lee, M.S., Ryu, S.H., Miyazawa, K., Park, S.H., Cho, C.S. and Kim, W.U. (2006) ‘Calcineurin is expressed and plays a critical role in inflammatory arthritis’, J Immunol, 177(4), 2681–90, available: http://dx.doi.org/10.4049/jimmunol.177.4.2681.

Zerbino, D.R., Achuthan, P., Akanni, W., Amode, M.R., Barrell, D., Bhai, J., Billis, K., Cummins, C., Gall, A., Giron, C.G., Gil, L., Gordon, L., Haggerty, L., Haskell, E., Hourlier, T., Izuogu, O.G., Janacek, S.H., Juettemann, T., To, J.K., Laird, M.R., Lavidas, I., Liu, Z., Loveland, J.E., Maurel, T., McLaren, W., Moore, B., Mudge, J., Murphy, D.N., Newman, V., Nuhn, M., Ogeh, D., Ong, C.K., Parker, A., Patricio, M., Riat, H.S., Schuilenburg, H., Sheppard, D., Sparrow, H., Taylor, K., Thormann, A., Vullo, A., Walts, B., Zadissa, A., Frankish, A., Hunt, S.E., Kostadima, M., Langridge, N., Martin, F.J., Muffato, M., Perry, E., Ruffier, M., Staines, D.M., Trevanion, S.J., Aken, B.L., Cunningham, F., Yates, A. and Flicek, P. (2018) ‘Ensembl 2018’, Nucleic Acids Res, 46(D1), D754–D761, available: http://dx.doi.org/10.1093/nar/gkx1098.

Zhou, Y. and Lingle, C.J. (2014) ‘Paxilline inhibits BK channels by an almost exclusively closed-channel block mechanism’, J Gen Physiol, 144(5), 415–40, available: http://dx.doi.org/10.1085/jgp.201411259.

